# Visual QC Protocol for FreeSurfer Cortical Parcellations from Anatomical MRI

**DOI:** 10.1101/2020.09.07.286807

**Authors:** Pradeep Reddy Raamana, Athena Theyers, Tharushan Selliah, Piali Bhati, Stephen R. Arnott, Stefanie Hassel, Nuwan D. Nanayakkara, Christopher J. M. Scott, Jacqueline Harris, Mojdeh Zamyadi, Raymond W. Lam, Roumen Milev, Daniel J. Müller, Susan Rotzinger, Benicio N. Frey, Sidney H Kennedy, Sandra E. Black, Anthony Lang, Mario Masellis, Sean Symons, Robert Bartha, Glenda M MacQueen, The CAN-BIND Investigator Team, The ONDRI Investigators, Stephen C. Strother

**Author notes:** These authors contributed equally (Tharushan Selliah, Piali Bhati).

## Abstract

Quality control of morphometric neuroimaging data is essential to improve reproducibility. Owing to the complexity of neuroimaging data and subsequently the interpretation of their results, visual inspection by trained raters is the most reliable way to perform quality control. Here, we present a protocol for visual quality control of the anatomical accuracy of FreeSurfer parcellations, based on an easy-to-use open source tool called VisualQC. We comprehensively evaluate its utility in terms of error detection rate and inter-rater reliability on two large multi-site datasets, and discuss site differences in error patterns. This evaluation shows that VisualQC is a practically viable protocol for community adoption.

## Introduction

Morphometric analysis is central to much of neuroimaging research, as a structural T1-weighted magnetic resonance imaging (sMRI) scan is almost always acquired in all neuroimaging studies for a variety of reasons. The sMRI scans are used in a number of important ways including as a reference volume for multimodal alignment, delineating anatomical regions of interest (ROIs), and deriving a number of imaging markers such as volumetric, shape and topological properties. FreeSurfer (FS) is a popular software package for fully-automated processing of structural T1 weighted MRI (T1w-MRI) scans, often to produce whole-brain cortical reconstruction of the human brain, including the aforementioned outputs (Fischl 2012). Hence, rigorous quality control (QC) of FS outputs is crucial to ensure the quality, and to improve the reproducibility of subsequent neuroimaging research results.

FreeSurfer processing is often completed without any issues when the properties of input sMRI scans are favorable for automatic processing. The ideal characteristics of the input sMRI scans include, but are not limited to, strong tissue contrast, high signal-to-noise ratio (SNR), absence of intensity inhomogeneities, absence of imaging artefacts (e.g., due to motion and other challenges during acquisition) and lack of pathology-related confounds. In the absence of one or more of such ideal characteristics, which is often the case in large multi-site neuroimaging studies, and owing to the challenging nature of the fully automatic whole-brain reconstruction, FS processing leads to errors in parcellation. Failure to identify and/or correct such errors could result in inaccurate and irreproducible results. Hence, robust FS QC is crucial.

Research into developing assistive tools and protocols for the QC of morphological data can be roughly divided into the following categories:

- visual protocols for rating the quality of the sMRI scan as a whole (Backhausen et al. 2016; Marcus et al. 2013). These protocols are helpful as QC of *input* sMRI is required at the MRI acquisition stage (e.g., to increase sample sizes) as well as at the subsequent archival and sharing stages (to improve the quality and reproducibility of analyses)
- assistive tools (manual as well as automatic) to expedite the algorithms for automated assessment of the sMRI quality (Raamana 2018; SIG 2019; Woodard and Carley-Spencer 2006; Gedamu, Collins, and Arnold 2008; Rosen et al. 2017; Esteban et al. 2017; Keshavan et al. 2018; Klapwijk et al. 2019; White et al. 2018). Some of these tools may employ image quality metrics (IQMs) (Shehzad et al. 2015), or metrics from derived outputs produced by FreeSurfer and related tools, to aid in the prediction of scan quality. The IQMs can be extracted directly from the scan itself (e.g., properties of intensity distributions) or be based on one or more of the FreeSurfer outputs (e.g. Euler number, volumetric and thickness estimates etc)
- image processing algorithms to detect imaging artefacts such as motion, ghosting etc (Pizarro et al. 2016; Alfaro-Almagro et al. 2018; Mortamet et al. 2009).

However, much of the previous research has been limited to rating the quality of input sMRI scan, but not the quality of subsequently derived outputs such as FreeSurfer parcellation. The FreeSurfer team provides a <u>troubleshooting guide</u> (Freesurfer Team 2017), that is a series of visual checks and manual edits for a diverse set of the outputs it produces. While this guide is comprehensive, it is quite laborious to perform even for a single subject, presents a steep learning curve to typical neuroimaging researchers, and is simply infeasible to employ on the large datasets that are commonplace today. Hence, assistive tools and protocols to expedite or automate this tedious FS QC process are essential. There has been notable effort in developing protocols (ENIGMA Consortium 2017) as well as assistive tools (Keshavan et al. 2018; Klapwijk et al. 2019) for the QC of FreeSurfer outputs. While the mindcontrol webapp (Keshavan et al. 2018) is more accessible (being browser-based) and provides easy navigation through the dataset, the overall QC process is no different from the FreeSurfer’s recommended troubleshooting guide (which employs tkmedit and slice-by-slice review), and hence is still slow and labor-intensive. While operating in the cloud using a browser interface may present some benefits of accessibility, the complicated initial setup creates an additional barrier for non-expert users (large amounts of costly cloud storage space), issues related to privacy and anonymization (transferring imaging data to the cloud), as well as creating a major dependency on the cloud makes it unreliable and/or slow. Moreover, it does not present the important visualizations for pial surface (see Figure 1, Panel B), which are necessary to identify any topological defects.

**FIGURE 1:**
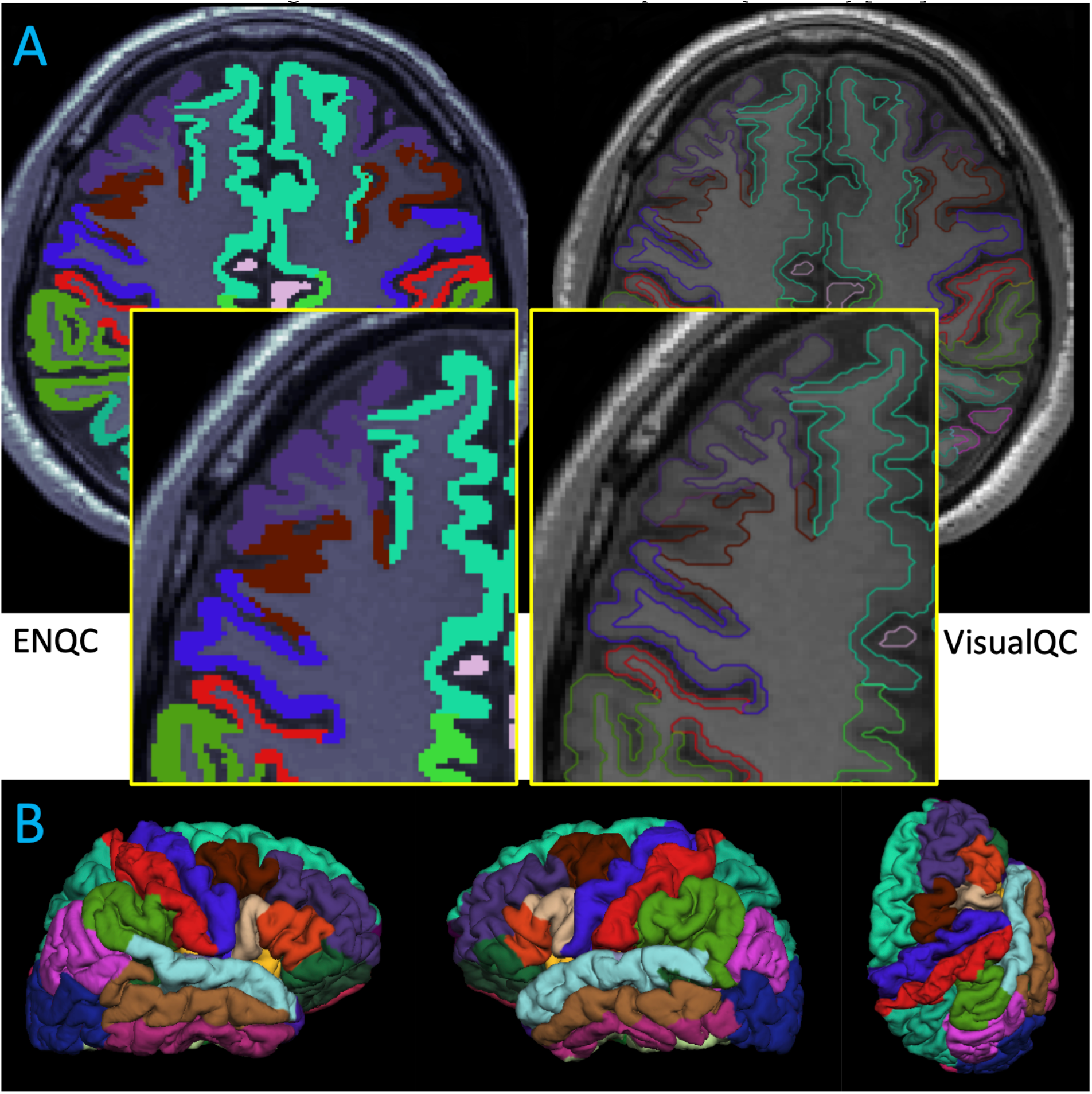
Panel **(A)**: Example illustrations of a single slice presented in the ENQC and VisualQC workflows respectively. The opaque overlay of cortical labels in ENQC makes it harder to see the boundaries of white and gray matter, and leads to errors in judging the accuracy of pial/white surfaces. Panel **(B)**: Illustration of external surface visualizations annotating a typical FS parcellation on the fsaverage surface. These are integrated into the default interface of VisualQC to enable easy detection of any topological defects and mislabellings, which is not the case with ENQC creating additional sources of error and burden.

In another related effort to reduce the QC burden as well as rater subjectivity, (Klapwijk et al. 2019) developed a statistical model to automatically predict a composite quality rating based on a combination of properties of input T1w MRI scan (presence of motion) and a few checks on the FS outputs. Their predictive model demonstrated very good performance (>80% accuracy; varying depending on evaluation setup) in discriminating “Failed scans” from the rest (rated as Excellent, Good or Doubtful). However, the rater agreement in this manual QC protocol was as low as 7.5% i.e. only six subjects out of 80 had ratings with a complete agreement among the five raters, increasing to >85% when majority rating is used to evaluate agreement. This may likely be due to the composite rating used (based on both input T1w MRI scan and FS outputs), which confounds the ratings, making it harder to disambiguate the source of bad quality (input vs. output), and hence making it a non-ideal comparison target. Moreover, their extensive analyses clearly highlight an important need of reliable and accurate ratings with high inter-rater reliability (IRR).

Aiming to deliver a quick method to QC FreeSurfer outputs from multiple large datasets, the Enhancing Neuro ImaGing through Meta-Analysis (ENIGMA) consortium (Thompson et al. 2020) developed a fast and useful visual rating protocol for FS QC (denoted by ENQC) based on a set of batch processing scripts, visualizations embedded in html and manual ratings collected in a spreadsheet. ENQC is a practical approach to greatly expedite an otherwise tedious process by selecting 4 volumetric slices for inspection. While drastically reducing the amount of work for the rater, this also greatly increases the likelihood of missing subtle errors, as they may fall between or outside the limited number of views. Moreover, having to deal with multiple disparate tools without clear integration (spreadsheets, shell scripts and html etc) leads to much higher human error (in maintaining integrity across multiple spreadsheets with complex identifiers), especially in large datasets.

To address the complexity and limitations of the various tools we mentioned so far (including ENQC) and the need for more reliable and accurate QC ratings, we developed VisualQC (Raamana 2018), a new open source QC rating framework, designed to ease the burden involving any visual QC tasks in neuroimaging research. The tool to rate the quality of FS parcellations is one of the many within VisualQC, which are built on a generic visual rating framework that is modular and extensible, allowing for manual/visual QC of virtually any digital medical data. Other tools within VisualQC include quality rating and artefact identification within T1w MRI, EPI and DTI scans, as well as tools to easily check the accuracy of registration, defacing, and volumetric segmentation algorithms. VisualQC’s custom-designed rating interface for FreeSurfer parcellation provides a seamless workflow, integrating all the necessary data and visualizations to achieve a high rating accuracy.

Based on a systematic study of two large multi-site datasets, from the Ontario Brain Institute (OBI): the Canadian Biomarker Integration Network in Depression (CAN-BIND) and the Ontario Neurodegeneration Research Initiative (ONDRI) programs, we show that the VisualQC protocol leads to a higher and more reliable error detection rate (EDR) than ENQC. As visual inspection is a subjective process, it is prone to bias or variation in a rater’s judgement and interpretation, especially in the case of subtle errors and those within tricky regions (with convoluted contours on 2D cross-sectional slices) such as entorhinal cortex, parahippocampal gyrus and superior temporal sulcus etc. Hence, we also quantify the inter-rater reliability (IRR) for each combination of dataset, for the two protocols ENQC and VisualQC. Our goal in choosing these two datasets is to evaluate the protocols on a diverse range of participants. In addition, we also chose to evaluate the QC protocols for two different versions of FreeSurfer: v5.3 and v6.0, as the parcellation accuracy and error patterns differ for different versions, and these two have been in use widely. These combinations would expose our study to diverse range of issues, as well as test the reproducibility and robustness of the protocol to differing datasets and software versions. Given the multi-site nature of these datasets, we also investigated site-wise differences in error patterns of FreeSurfer cortical parcellations. In particular, we built a predictive model of site to identify the factors influencing site-wise differences in FS error patterns. Based on this comprehensive evaluation, we show that VisualQC outperforms ENQC for FreeSurfer QC, becoming a strong candidate for a community consensus protocol for the visual QC rating of FS parcellations.

## Methods

### Datasets

We analyzed two large multi-site datasets that were drawn from previous studies 1) the Canadian Biomarker Integration Network in Depression (CAN-BIND) with 308 participants (MacQueen et al. 2019; Lam et al. 2016), and 2) the Parkinson’s disease cohort from the Ontario Neurodegeneration Research Initiative (ONDRI) (Farhan et al. 2017; Scott et al. 2020), with 140 participants. The demographics of the two datasets are shown in Table 1. More detailed information on site-differences, in terms of vendors, models and acquisition parameter information, is presented in Appendix A.

**TABLE 1:** Demographics for the two multi-site datasets in this study

| Dataset | N | Male/Female | Age | Group |
| --- | --- | --- | --- | --- |
| CANBIND | 308 | 110/198 | 34.45 (12.13) | Healthy controls (n=111)<br>Major depressive disorder (n=197) |
| ONDRI | 140 | 109/31 | 67.94 (6.35) | Parkinson's Disease (n=140) |

All statistics here are displayed in mean (SD) format.

### Processing

All scans in the two datasets were processed with the FreeSurfer cross-sectional pipeline (Fischl 2012), to obtain the default whole-brain reconstruction with no special flags. No manual editing was performed on the output parcellation from FreeSurfer, to focus the analysis purely on fully automatic processing. Each dataset was processed with two widely-used versions of 5.3 and 6.0, on a CentOS 6 Linux operating system in a Compute Canada high-performance computing cluster.

### Rating Methodology

The primary purpose of FS QC via manual visual rating is to identify parcellation errors and rate their level e.g. as Pass, Major error, Minor error, [complete] Fail etc. An error in FS cortical parcellation occurs when the pial or white surface do not follow their respective tissue-class boundaries, such as gray matter (GM) and white matter (WM) respectively.

Initially error inspection was completed by three raters, following protocols from ENQC. Briefly, ENQC rates the quality of the parcellation based on two types of visualizations: 1) *Internal QC*: four cross-sectional slices in coronal and axial views overlaying the labels voxel-wise on top of the input T1w MRI in opaque color (see Figure 1), and 2) *External* QC: four views of the anatomical regional labels visualized on the *fsaverage* surface^1^. If there are no issues of any kind in the internal or external QC, it is rated as Pass in that corresponding section. Parcellation errors localized to particular regions are labelled as Moderate, whereas presence of severe errors, large mislabeling, mis-registration and imaging artefacts as well as global failures would be rated as Fail. Location of the error, in terms of left (L) or right (R) hemispheres as well as the particular region of interest (ROI), are also noted, following the FreeSurfer Color Lookup Table (FSCLUT) [<u>link</u>].

The FS QC interface for VisualQC is shown in Figure 2. This is highly customized for rating the accuracy of FS parcellation, and presents a comprehensive picture in all the relevant views: contours of pial and white surfaces in all three cross-sectional views with at least 12 slices per view (default is two rows of six slices, but it is customizable), along with six views of the pial surface (in the top row). The cortical labels in both the cross-sectional and surface views are color-annotated in the same manner as the Freesurfer’s *tksurfer* tool to leverage the familiarity of the default color scheme. This design, while rigorous, still allows for rapid review of the quality and bookkeeping of the rating along with any other notes.

**Figure 2:**
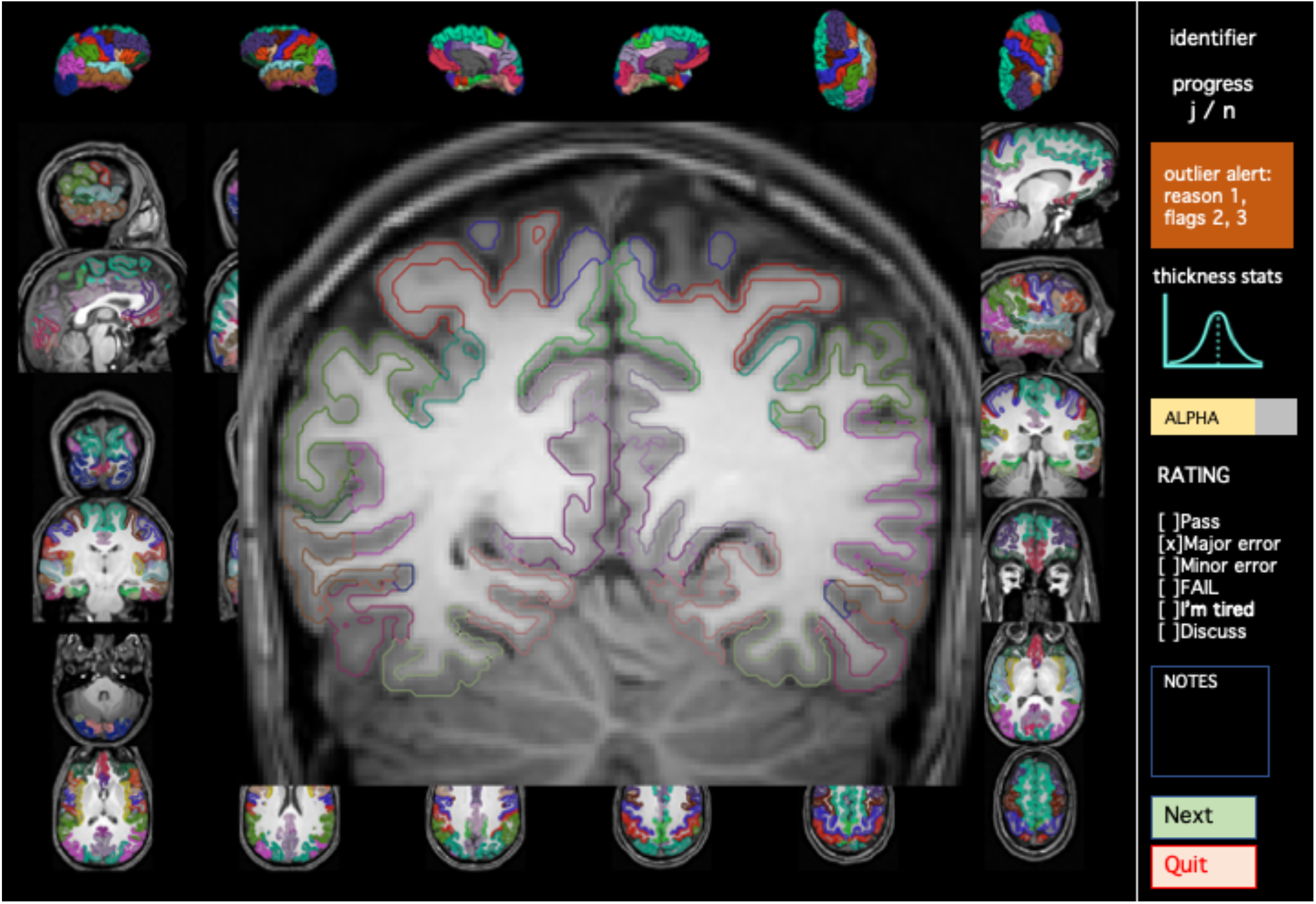
An instance of the VisualQC interface for the rating of parcellation accuracy of FreeSurfer output. This customized interface presents a comprehensive picture of the parcellation in all the relevant views: contours of pial and white surfaces in all three cross-sectional views with 12 slices each, along with six views of the pial surface, color-annotated with corresponding cortical labels. This design, while rigorous, still allows for rapid review of the quality and bookkeeping of the rating along with any other notes.

For our three raters, compared to ENQC, VisualQC enabled recording additional intermediate levels, encoded as *Pass, Minor Error, Major Error* and [complete] *Fail*. The locations of parcellation errors are also noted in VisualQC using the Notes section in the rating interface below the radio buttons for rating, using the same names and codes as in the FSCLUT.

FreeSurfer parcellation errors can be roughly categorized as in Table 2 below. Their names are self-explanatory, and their frequencies for these common errors are estimated from the ratings data presented in this manuscript. The detailed rating system, along with definitions and examples for different levels of errors is presented in Section 3.4 of the VisualQC manual at github.com/raamana/visualqc. The direct URL for the current version `v1.4` of manual is https://github.com/raamana/visualqc/blob/master/docs/VisualQC_TrainingManual_v1p4.pdf.

**Table 2:**
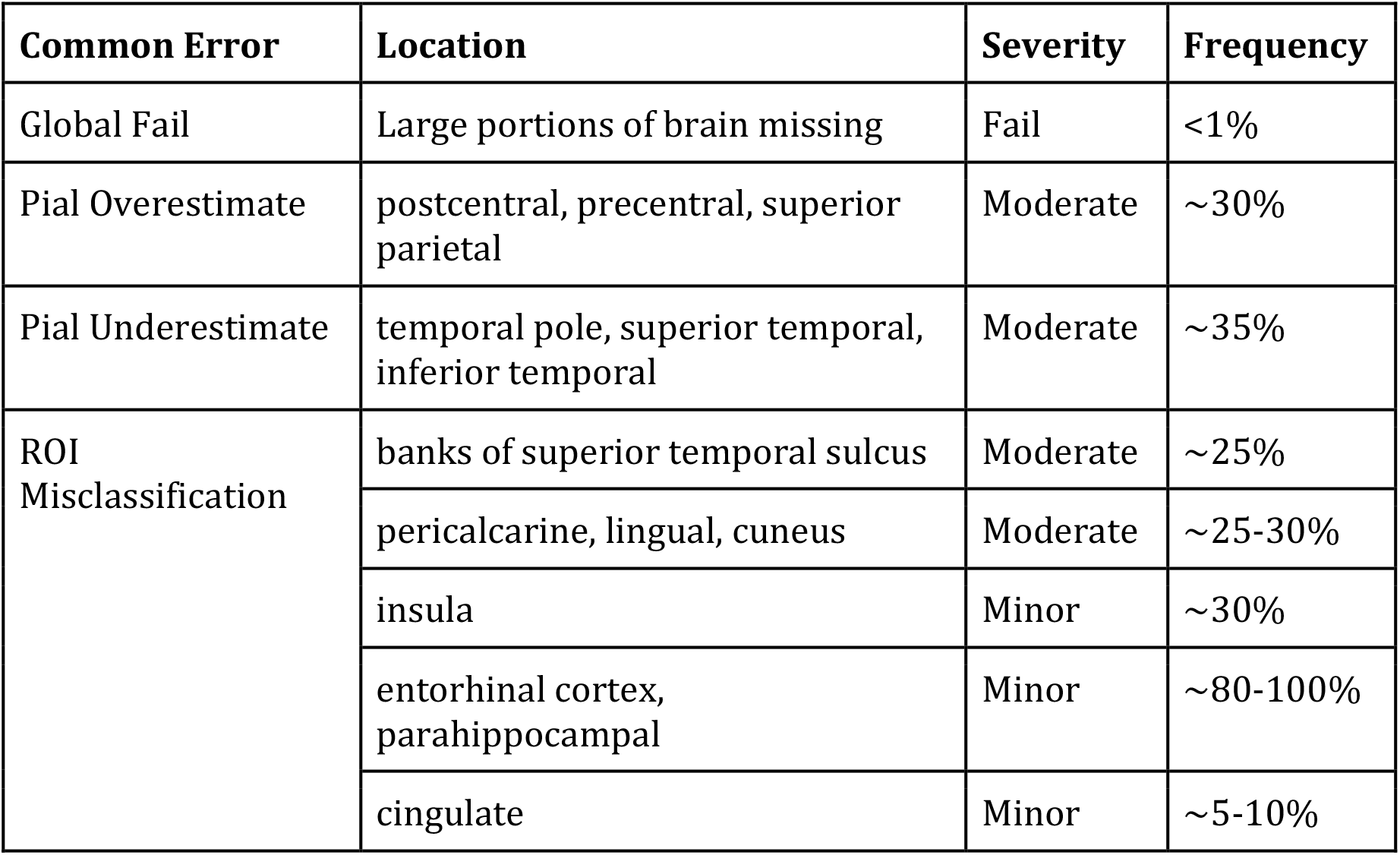
Rough categorization of the common parcellation errors from Freesurfer, their locations and frequencies.

### Exceptions to Rating

Accurate parcellation in highly convoluted areas such as the entorhinal cortex (EC), parahippocampal gyrus (PH) and insula (IN) is highly challenging. Although FreeSurfer is generally accurate in many regions of the cortex in the absence of imaging quality issues, it routinely is erroneous in these ROIs (see Figure 3, and quantification below). Minor errors in these ROIs are so common, ENQC protocol chose to rate them as *Pass* (ignoring them for the overall quality for the whole brain parcellation), so long as the errors are minor and the parcellation is free from any other issues. This is in line with the official troubleshooting guide (Freesurfer Team 2017) which recommends avoiding editing these minor errors to avoid introducing bias and reducing reliability. In the VisualQC protocol, we choose to note them as *Minor Error* in the interest of recording the most accurate reflection of the parcellation quality. Our data confirms these errors are almost universal: only 4/2688 ratings from three raters (0.1%) were free from errors in EC, PH and IN.

**FIGURE 3:**
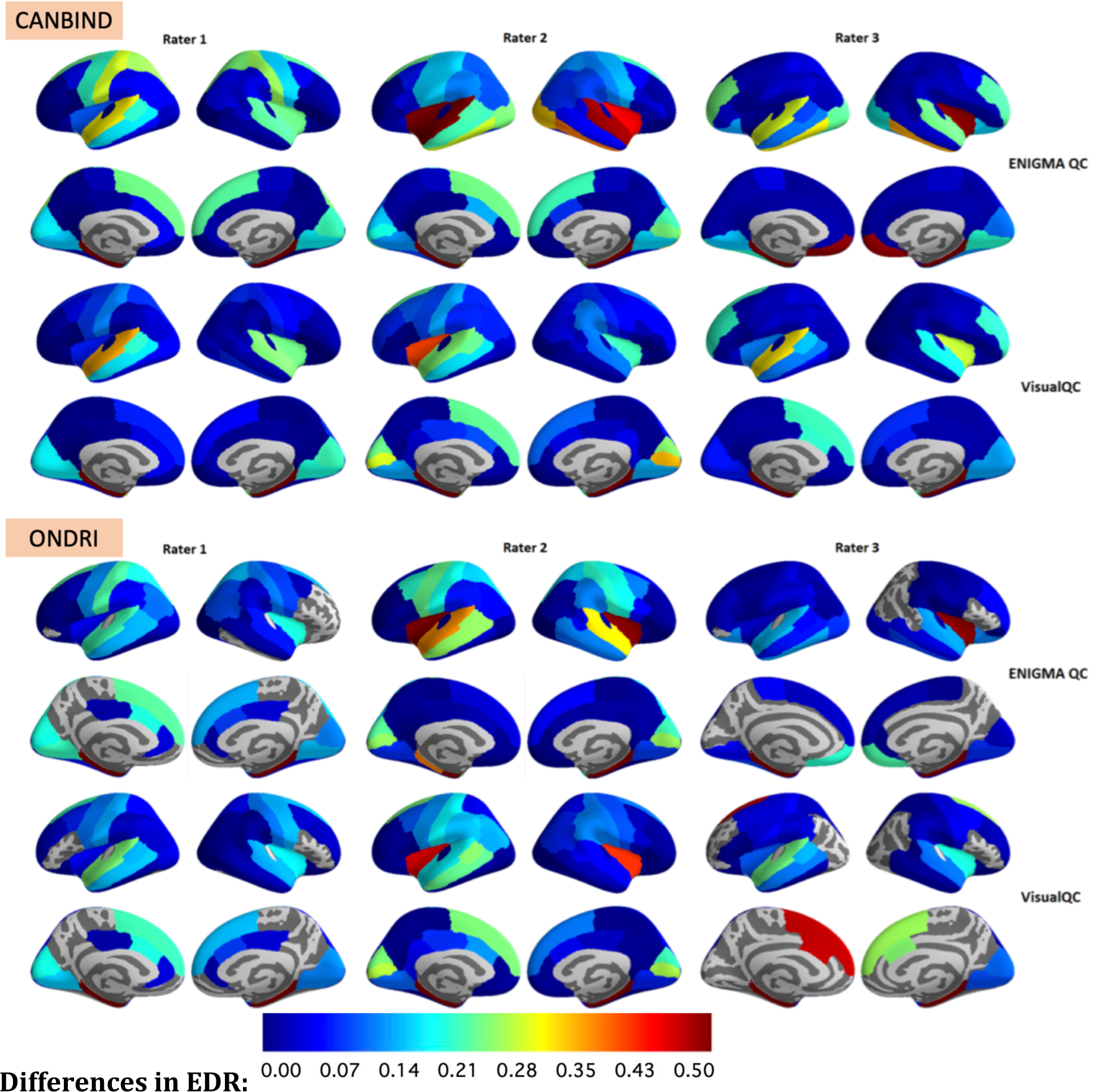
Visualization showing the differences in EDR across multiple raters for FreeSurfer v6.0 parcellations in the CANBIND and ONDRI datasets for ENQC and VisualQC protocols. All the visualizations in this paper are annotated with the default Desikan-Killiany parcellation unless otherwise stated. The non-coloured areas in gray are regions without any parcellation errors or where there is no cortex present (e.g. corpus callosum).

In our statistical analyses comparing error frequencies, we have recoded minor errors in EC, PH and IN with no other issues in VisualQC ratings as *Pass*, to make them commensurable with ENQC. A similar approach is taken with minor errors (over- and underestimates) in superior frontal (interacting with the cingulate gyrus), superior parietal (interacting with cuneus and/or precuneus), supramarginal gyrus (also impacts superior temporal (ST)), and middle temporal (MT) gyrus (interacting with inferior temporal (IT)).

### Error statistics

#### Error detection rate

Error detection rate (EDR) for a brain region was calculated as the number of participants with detected errors, divided by the total number of participants in that dataset. For valid comparison with VisualQC in quantifying EDR, we considered a parcellation as *Pass* in ENQC only when it is rated as *Pass* in both Internal and External evaluations, and as *Fail* for all other combinations. Under the VisualQC protocol, only *Pass* is considered *Pass* and any other rating as *Fail*. This statistic helps us judge which FS version is generally more accurate, and how that performance is related to experimental conditions (e.g. site, scanner). EDR was calculated separately for each dataset, FreeSurfer version, and rating protocol.

#### Inter-rater reliability

The ratings were hierarchical in nature as each rating was initially approached with a *Fail* vs. *Pass* mindset, which was then followed by dividing the *Fail* category further into multiple levels (Major vs. Minor vs. complete Fail). As the interval between the Major vs. Minor errors and Minor vs. Fail can and does differ, they cannot be treated as simple numerical variables. Given subjectivity in rating error severity - what one rater may perceive as minor error could be perceived as major error by another reviewer, esp. when traversing across the entire brain covering diverse ROIs, the ordering of error severities is not preserved across raters, and hence they cannot treated as ordinal variables either.

Therefore, we encoded them as categorical variables to produce valid statistics to respect their properties and measurement methods. Inter-rater reliability (IRR) for ratings was computed based on the most native form of ratings possible, such as “*Pass*”, “Major Error”, “Minor Error”, “Fail” for VisualQC. For ENQC, the concatenated ratings from Internal and External QC used for IRR calculations are “*Pass_Pass*”, “*Pass_Fail*”, “*Fail_Pass*” and “*Fail_Fail*”.

We quantified IRR using the Fleiss Kappa statistic on ratings from the three raters (Fleiss 1971; Randolph 2005), separately for each dataset, FreeSurfer version, and rating protocol. In addition, we have also bootstrapped this computation 100 times selecting 80% of the sample for each combination, to analyze the stability of estimates.

### Automatic Site Identification

Another way to demonstrate the site differences is by trying to automatically predict the site based on morphometric features, as they play a direct role in tissue contrast and hence FS accuracy. Towards this, we computed region-wise descriptive statistics (such as mean, SEM, kurtosis, skew, and range) on all cortical features (i.e., thickness, area, curvature) and contrast-to-noise ratio (CNR)^2^ values in all FS labels.

For site-identification, a random forest classifier was trained on the aforementioned features to predict the site label. We evaluated its performance with *neuropredict* (Raamana 2017; Raamana and Strother 2017) using repeated-holdout cross-validation (80% training, repeated 30 times; feature selection based on f-value). To clarify, this analysis is performed to demonstrate the presence of large site/scanner-differences in tissue contrast patterns as they play a key role in FS parcellation accuracy. This may not be directly related to the central goal of this paper i.e., evaluate and compare the visual QC protocols for Freesurfer. However, it tests our expectation that the primary site/scanner specific differences will be driven by basic CNR effects, and not other derived measures.

### Software

All calculations were performed based on the scientific Python ecosystem (Python version 3.6), with the Fleiss Kappa implementation coming from the `statsmodels` package version 0.10.1 (Seabold and Perktold 2010).

VisualQC is an open source QC rating framework (Raamana 2018) freely and publicly available at https://github.com/raamana/visualqc. The tool to rate the quality of FS parcellations is one of the many within VisualQC, which are built on a generic visual rating framework that is modular and extensible, allowing for manual/visual QC of virtually any digital medical data. Other tools within VisualQC include quality rating and artefact identification within T1w MRI, EPI and DTI scans, as well as tools to easily check the accuracy of registration, defacing and volumetric segmentation algorithms. They are documented in detail at https://raamana.github.io/visualqc/, which also includes a comprehensive manual to train the rater to learn and use VisualQC^3^.

## Results

### Error detection rate

The EDR measured by different raters in the CANBIND and ONDRI datasets for FS v6 are shown in Figure 3, which reveals the following: 1) there are some ROIs that are consistently picked up as erroneous by all raters using both QC packages, e.g. in the medial temporal lobe (MTL), such as the ET, ST and PH. This is not surprising given the challenges involved in producing an accurate parcellation in these challenging areas in a fully automatic fashion; 2) beyond the MTL, we notice variability in EDR patterns across the three raters, both between the two protocols, and even within the same protocol; 3) There is clear variability in EDR per region either across the raters within the same protocol, or across the protocols for the same rater. This is only to be expected given the subjective task across human raters. The regions where this variability is large, both across raters and protocols, are the hard-to-segment temporal lobe ROIs as well as the central sulcus.

### Error Comparison

Differences in EDR found between VisualQC and ENQC, computed as EDR(VisualQC)- EDR(ENQC) are shown in Figure 4, on the default Desikan-Killiany parcellation. We observe some interesting patterns in the difference plot. The majority of those differences in EDR can be divided into two categories:

**FIGURE 4:**
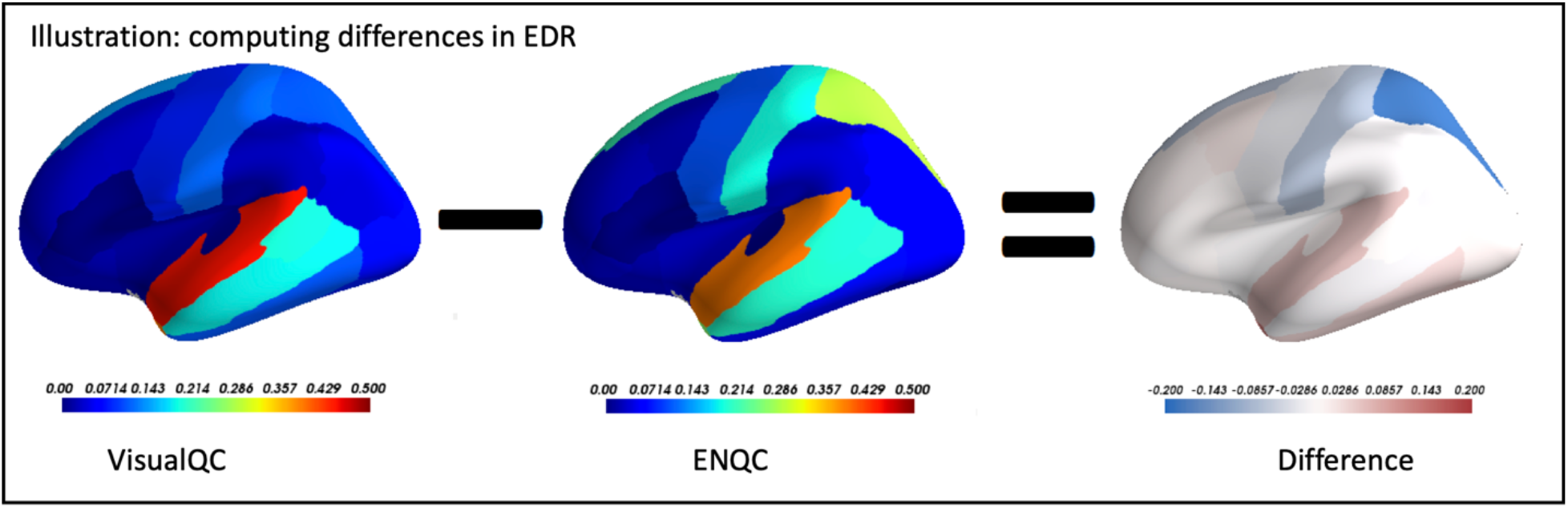
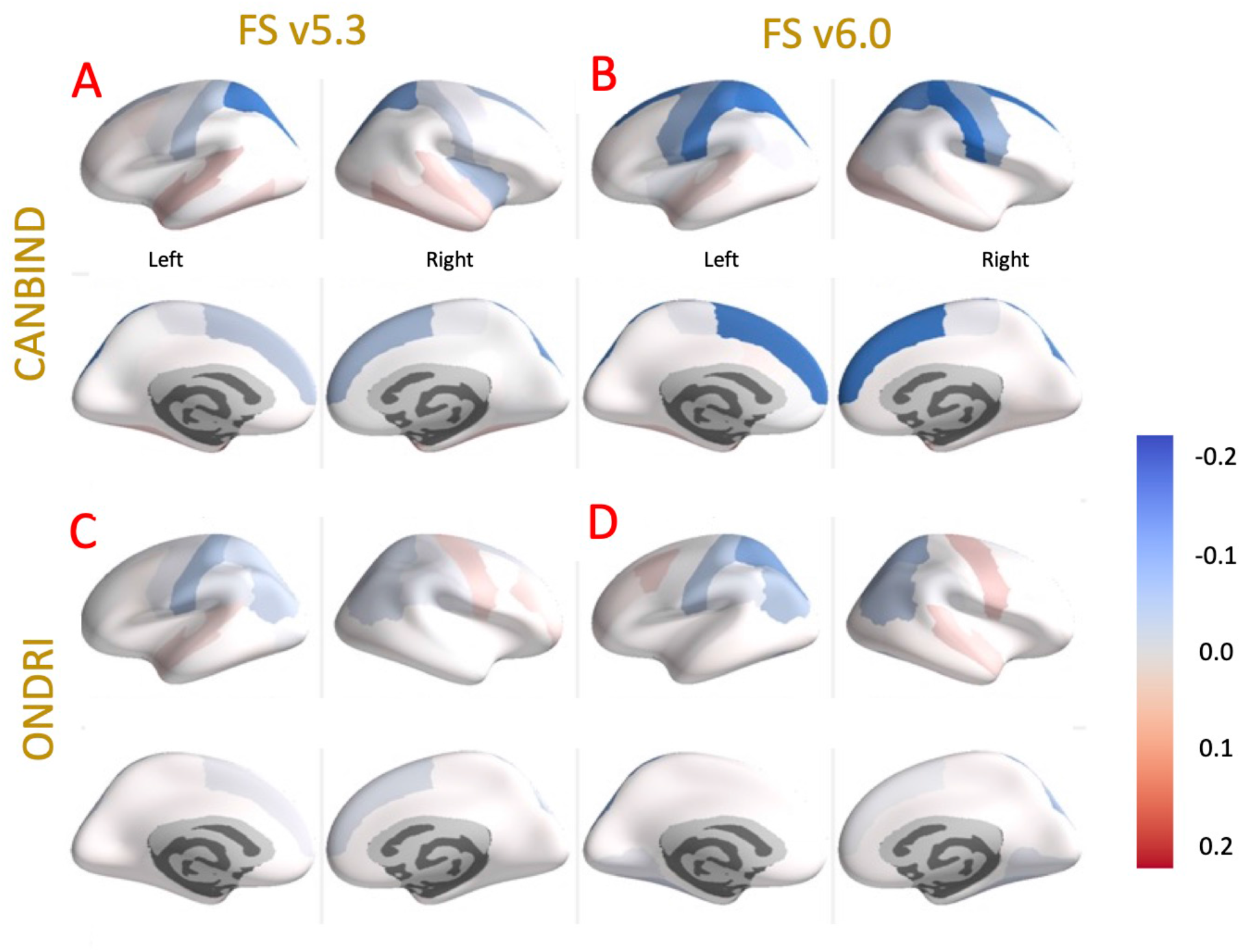
Percentage differences of error detection found between ENQC and VisualQC, where negative value (in blue) indicates that ENQC detected a greater percentage of errors, whereas a positive value (in red) indicates that VisualQC found greater percentage of errors, for that dataset and version of Freesurfer. The color bars for all panels visualizing the EDR differences range from -0.2 to 0.2. The four panels shown below are: **(A)** CAN-BIND, FS v5.3, **(B)** CAN-BIND, FS v6.0, **(C)** ONDRI, FS v5.3 and **(D)** ONDRI, FS v6.0. Each panel shows lateral/medial views of the EDR map in top/bottom rows respectively.

- a higher percentage of errors detected in the temporal poles by VisualQC, in slices below that of the lowest available view using ENQC, and
- a higher percentage of errors detected by ENQC in the upper pial surface (superior parietal lobule, superior frontal, pre- and postcentral sulcus), primarily in the CANBIND cohort.

Due to ENQC’s choice of an opaque overlay of segmentation labels onto the anatomical MRI (see Figure 1), this increased rate of error detection is likely due to a reduction in visibility of the structural scan itself, resulting in a higher false positive rate (FPR). Note: we believe errors identified via VisualQC are inherently more accurate by virtue of its superior design (much expanded coverage of the parcellation/brain, non-opaque contour overlay with the ability to tweak their transparency levels, including switching them off etc).

### Inter-rater reliability

The IRR estimates for different combinations of datasets and FreeSurfer versions are presented in Table 3 for the two protocols. This shows that VisualQC is more reliable across the board. In addition, the bootstrapped estimates (presented in Appendix B) are quite identical to those shown in Table 3. We believe this is due to presenting the rater with a vastly more comprehensive view of parcellation, the ability to zoom-in to each slice as well as toggle the tissue contour overlay to evaluate the anatomical accuracy in a confident manner.

**TABLE 3:** Inter-rater reliability (IRR) estimates for the three raters for different combinations of the dataset and FreeSurfer versions.

|  | CANBIND v6.0 | ONDRI v6.0 | CANBIND v5.3 | ONDRI v5.3 |
| --- | --- | --- | --- | --- |
| ENQC | 0.28 | 0.215 | 0.36 | 0.25 |
| VisualQC | 0.64 | 0.54 | 0.58 | 0.56 |

### Site differences

Given FS performance is dependent on the quality of the input T1w MRI scan and the underlying tissue contrast, we wanted to study if the acquisition site played any role in EDR and whether different sites presented different error patterns. Hence, we visualized the parcellation errors segregated by site, which are presented in Fig. 5 for the CANBIND dataset processed with FS v6.0. This visualization illustrates the large variability across sites in multiple ROIs of the brain across the cortex. This variability can also be observed even in the frequently erroneous temporal lobe regions.

**FIGURE 5:**
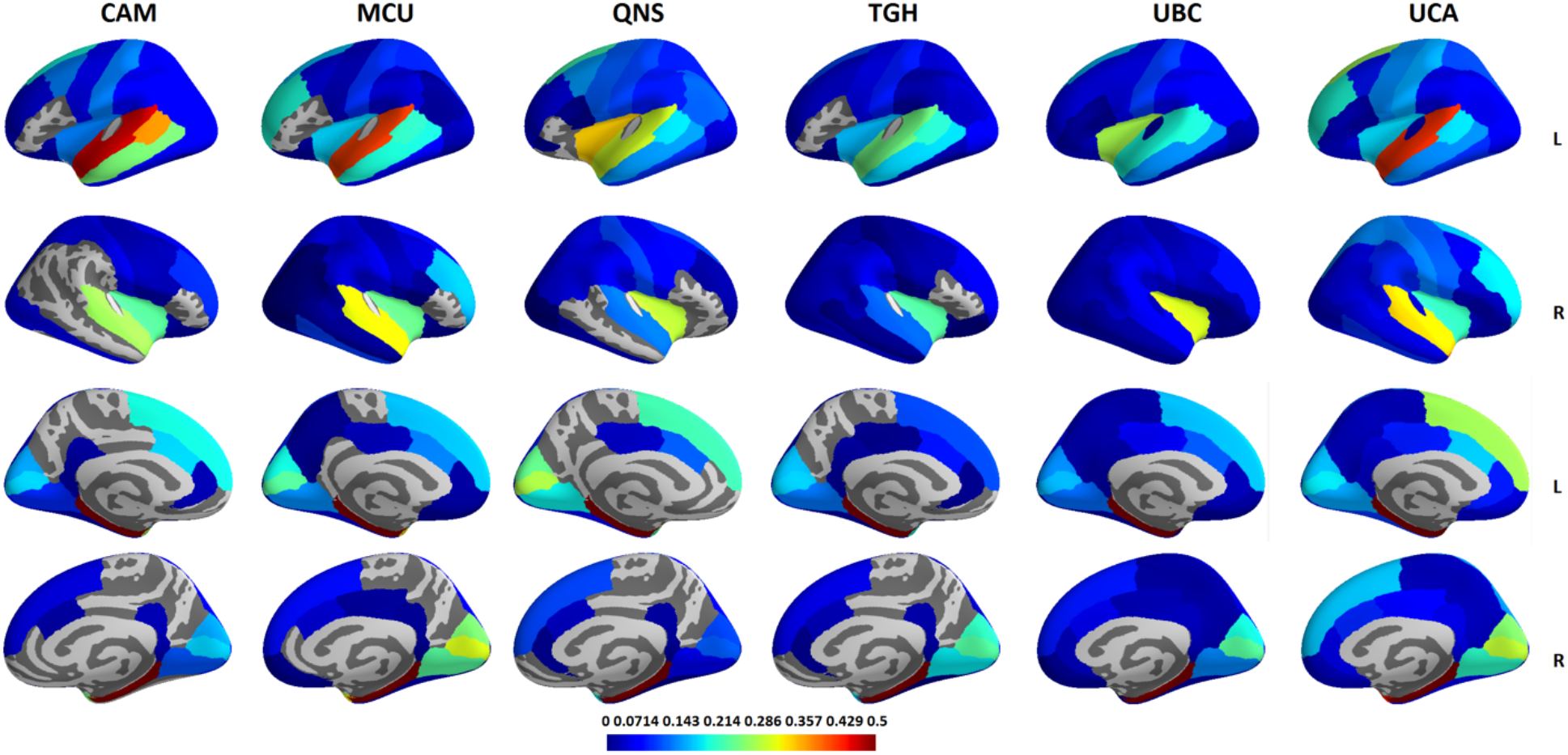
Visualization of the site differences in error ratings (average of the percent errors across the three raters) across different sites for the CANBIND dataset (FS v6.0)

The corresponding site differences for the ONDRI dataset (FS v6.0) are shown in Figure 6. We observe some clear patterns common across the sites here, such as the relatively higher error rate observed in the medial temporal lobe (MTL) and superior frontal (SF) cortex.

**FIGURE 6:**
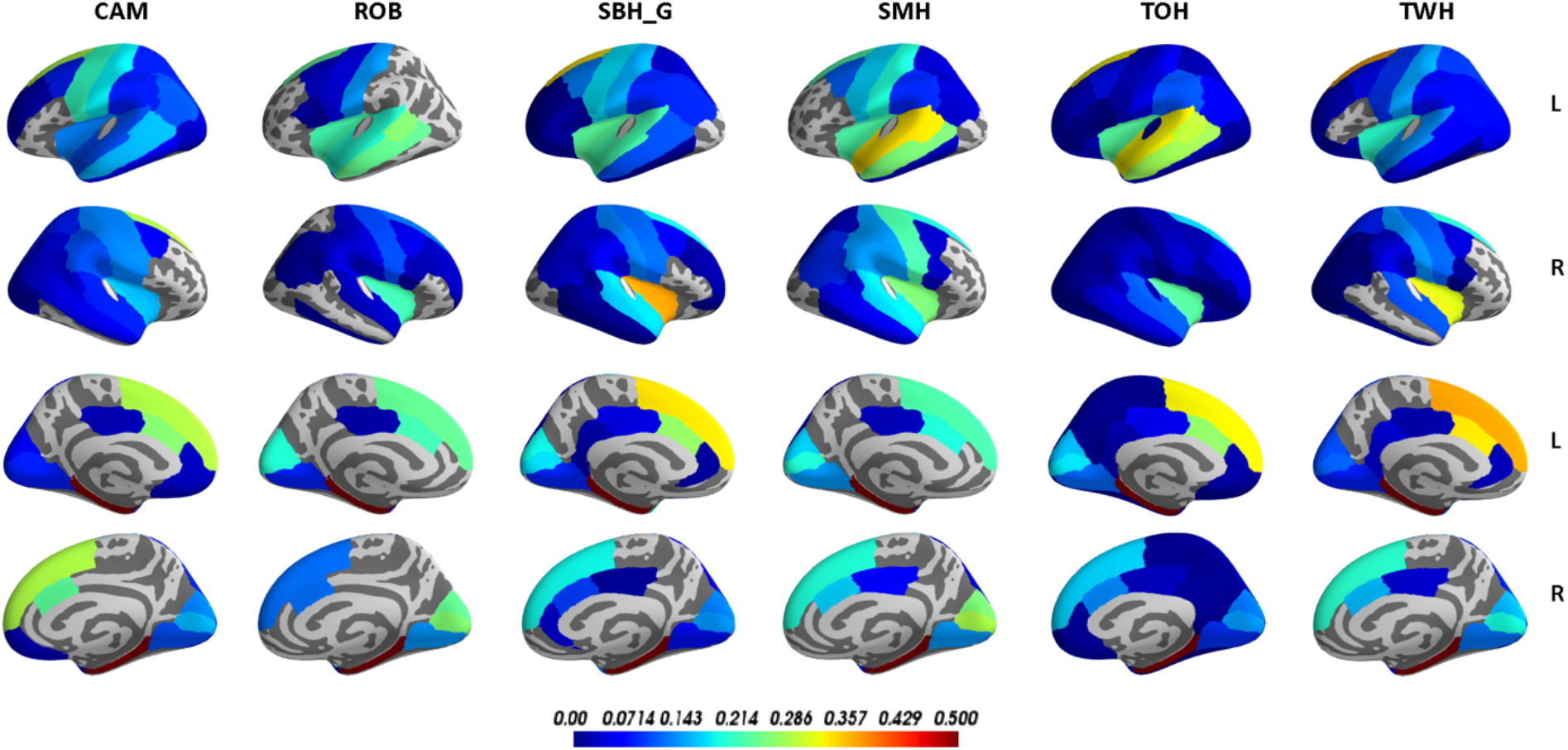
Visualization of the site differences in error ratings (average of the percent errors across the three raters) across different sites for the ONDRI dataset (FS v6.0)

Although higher error rate is expected in MTL, which was also observed in the CANBIND dataset, similar high error rate in SF is an interesting surprise.

### Automatic Site Identification

The performance estimates of a predictive model for automatic site identification on the FS v6 outputs from the CANBIND dataset are visualized in the confusion matrix shown in Figure 7. This shows some sites, especially UBC and QNS, are readily identifiable with over 80% accuracy. Given the chance accuracy in this 6-class experiment is 16%, sites TGH, MCU and UCA seem relatively easily identifiable as well.

**Figure 7:**
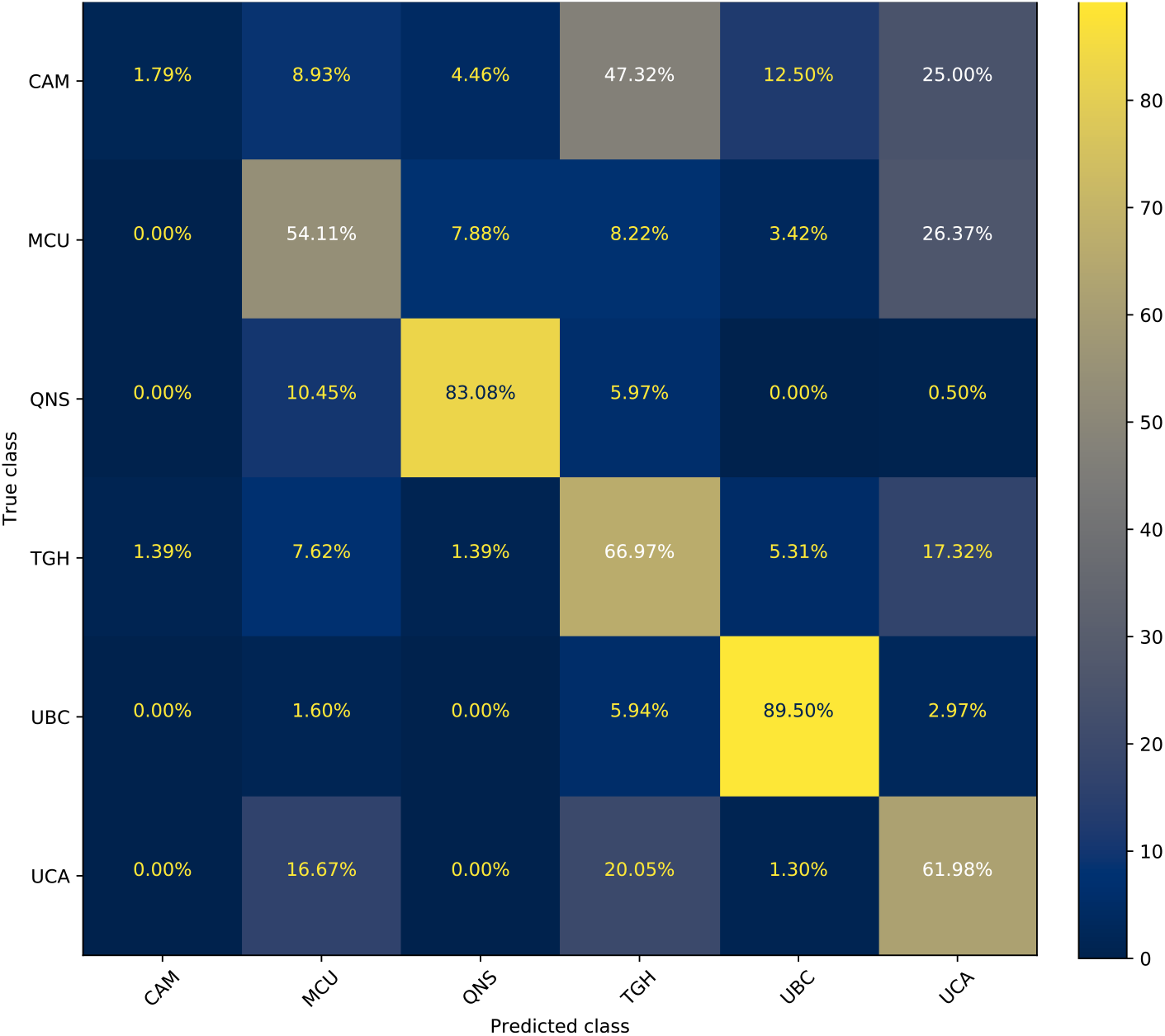
Confusion matrix from a simple machine learning experiment to identify site from the morphometric features extracted from FreeSurfer outputs (v6.0) from the CANBIND dataset, such as the region-wise statistics on all cortical features (thickness, area, curvature) and CNR values in the FS labels. We notice some sites, esp. UBC and QNS, are quite identifiable. Given the chance accuracy in this 6-class experiment is 16%, sites TGH, MCU and UCA seem easily identifiable also.

It is rather interesting CAM and MCU have often been misclassified (>25%) as UCA, which can also be seen in the similarity of site-wise error patterns in Figure 5. Moreover, all these 3 sites use the same scanner (GE 3.0T Discovery MR750), which might explain the confusion exhibited by the site-predicting-classifier. However, it must be noted CAM also got misclassified as TGH 47% of the time, whereas TGH has never been misclassified as CAM (1.39%). Such asymmetric prediction might have been a result of small sample size for CAM (n=16), which might be causing challenges for the predictive model in learning a unique profile for this site and/or skew towards majority classes to improve performance. This anomaly is interesting and worthy of further future investigation. Please refer to Appendix A for more details on the scanner models and acquisition parameters.

The corresponding feature importance values (median values from the 30 repetitions of cross-validation) are visualized in Figure 8. It is quite clear from the top 10 features that CNR played a crucial role in site identification, and their source ROIs are in challenging areas such as the lateral occipital cortex, fusiform gyrus, cuneus, postcentral gyrus, superior parietal cortex and temporal lobe. These site-differentiating ROIs are difficult to identify just based on raw patterns shown in visualizations such as Figure 5.

**Figure 8:**
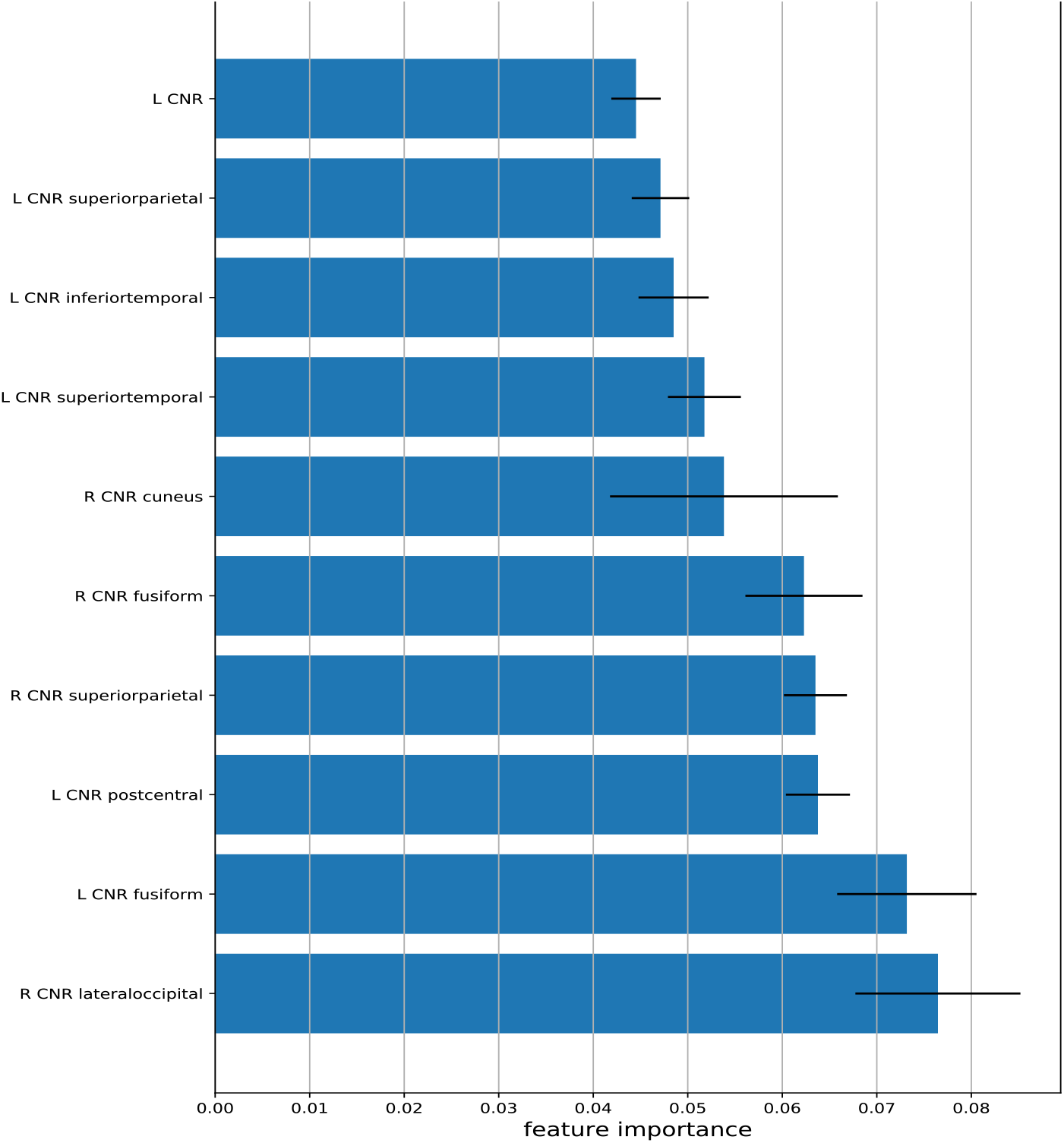
Feature importance values from the random forest predictive model for site identification on CANBIND dataset FS v6.0.

We have also performed the site differences analysis on the ONDRI dataset (FS v6.0) with results shown in Figure 9. Similar to CANBIND, we can see that a few sites are quite identifiable in ONDRI as well, such as TOH and TWH with 84% and 71% accuracy. Given the chance accuracy in this 5-class experiment is 20%, we can consider the sites LHS and SBH to be identifiable as well. The features contributing most to the automatic site identification model were sulcal depth in rostral anterior cingulate and precentral gyrus, thickness distributional statistics (such as mean, skew, range, SEM) in paracentral, inferior temporal, lingual and precentral gyri, along with precuneus volume (fraction relative to the whole brain). It is interesting to note these features are a different set compared to those in CANBIND which were mostly based on CNR profiles in different ROIs. Although the site prediction analyses presented here are based on derived features, and error patterns across sites are based on raw parcellations of white and gray matter surfaces, the site-prediction results from the two datasets show the importance of being cognizant about site differences while QCing FS parcellations.

**Figure 9:**
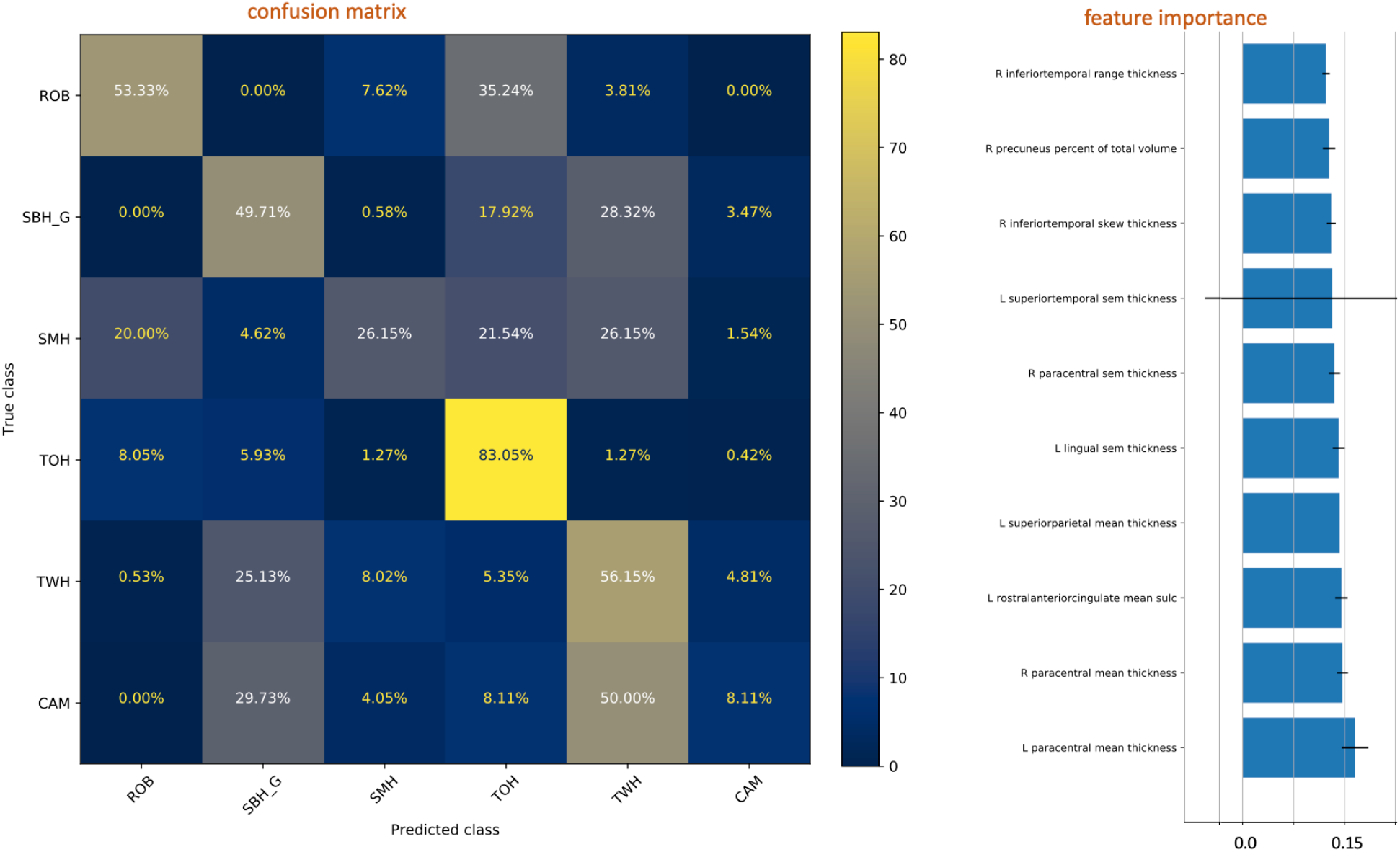
Confusion matrix (left panel) from of the predictive model for site identification based on FS outputs (v6.0) from the ONDRI dataset. We utilize the same features as were extracted in the CANBIND dataset. The corresponding feature importance values are shown in the right panel.

## Discussion

In this paper, we visualize the patterns in EDR for two different external protocols and evaluate their utility and reliability in different dimensions. In the context of FS QC broadly speaking, the Troubleshooting guide (TSG) recommended by the Freesurfer team is another common occurrence. TSG is a useful practical guide to not just identify common issues but also fix them with specific changes to intermediate outputs and with some additional processing. However, it is not a quality rating protocol per se, like ENQC or VisualQC, as it is a guide towards identifying common errors Freesurfer makes and how to fix them via manual editing. The approach recommended in the TSG boils down to traversing every single slice one at a time and checking the parcellation accuracy, which although being close to the best one can do (*gold standard*), is quite time consuming and simply not feasible for even for somewhat small datasets, let alone large datasets. That’s the basis for the development of protocols like ENQC and VisualQC. As for the EDR, following the TSG would result in labeling almost all the parcellations as erroneous. Based on our experience of QCing 1000s of scans from many datasets covering a gamut of demographics and sites, we are confident the EDR would be 100% when following the TSG or any other process requiring inspection of every single slice/ROI. EDR would be very close to 100% not just because Freesurfer algorithms have issues but mainly due to the immense complexity involved in the whole brain reconstruction process in a fully automatic fashion. As noted in the Methods sub-section “Exceptions to Rating”, the EDR for VisualQC taking all the minor errors identified into account, is 99.9% (only 4 combinations out of 2688 were free from any errors). To reinforce the point, this was based on only looking at the 36 cross-sections, and if we increased them further, we would very likely identify issues in those 4 combinations as well. This implies 1) there were no false positives and 2) there is no loss of quality of rating in employing VisualQC protocol, and there are only benefits to be had in terms of efficiency and productivity.

In addition to rating accuracy, protocol efficiency is important given the steadily increasing size of the neuroimaging datasets. Relative to VisualQC, ENQC is slower and more erroneous because of need to obtain and manage a disparate collection of tools (n>3: external QC, internal QC, separate spreadsheets for note taking, outlier prompts etc) to rate a single subject, whereas that is all fully integrated and seamless in a single VisualQC interface. We estimate our interface would be roughly at least 3-5 times faster. In the default configurations, we estimate it takes about 20 secs for a trained rater in VisualQC whereas it may take a minute or two in ENQC. That is reinforced further when we take the ease of initial installation and future upgrades into account (one command for us vs. manual management of many for ENQC). That is reinforced further when we take the ease of initial installation and future upgrades into account (one command for us vs. manual management of many for ENQC). It must be noted the efficiency/processing times can vary based on the type of configuration one may choose for VisualQC and goals of the specific QC task (# slices per view, series and type of checks made, along with how thorough is with the notes they make).

While we find the IRR for VisualQC is relatively higher than ENQC, we can further improve it in a few ways e.g. by reducing the subjectivity in the rating system when possible. Discounting the irreducible human subjectivity, we can design the training protocol to be more comprehensive to develop consensus on typical disagreements. Another possibility could be to increase the number of checkpoints to review before rating, but this option comes with the tradeoffs of additional burden and slower processing time.

As easy and integrated as VisualQC is, manual QC still is not effortless, especially with the increasingly large sample sizes reaching many 10s of thousands today. Hence, an automated tool to predict the quality of a given FS parcellation without human rating would be useful in reducing the QC burden. A frequently requested feature is an automatic tool to identify clear failures and major errors sufficiently accurately, so the raters can focus on the subtle and minor errors, which would expedite the QC process significantly.

However, as highlighted by previous efforts in this direction (Klapwijk et al. 2019), the development of accurate automatic predictive QC tools require that we have a reliable approach to create ground truth (via visual QC) for these tools to be trained on and optimized for. Development of such a reliable protocol as a candidate for community adoption was the main thrust of this paper. Based on this protocol, we plan to pursue to development of a predictive tool and validate it for different application scenarios such as high sensitivity (not missing even a single bad parcellation) or more narrowly to clear certain ROIs (posterior cingulate gyrus or medial temporal lobe etc) of any errors. Other frequently requested features that VisualQC does not currently support, but plan to develop in the future, are 1) the ability to *correct* the errors as they are identified on the VisualQC interface, 2) automatic recording of the location(s) of the erroneous parcellation, 3) ability to dig deeper (via zooming in or selective highlights) on a single ROI (such as precuneus) while switching off everything, and 4) intelligent slice selection or incorporating application-specific domain knowledge to improve the speed or accuracy of the visual QC task at hand.

## Conclusions

In this study, we presented a viable protocol for the visual QC of FreeSurfer parcellations based on an open source QC library. Based on a systematic comparison, we demonstrate that this VisualQC protocol leads to relatively higher EDR, lower FPR and higher IRR for the manual QC of FreeSurfer parcellation relative to ENQC. We characterized its utility and performance on two large multi-site datasets showing it is robust across two different age ranges and disease classes. Moreover, it is seamless and is significantly faster than following ENQC or the standard FreeSurfer troubleshooting guide. Further, we highlight the need to be cognizant of the site-differences in parcellation errors.

## Appendix A Site information

The two datasets studied here are large and multi-site by design. The detailed information on site-differences in terms of acquisition parameters and scanners have been carefully tabulated in the respective dataset papers for ONDRI (Scott et al. 2020) and CANBIND (MacQueen et al. 2019).

CANBIND:
| Site | Toronto General/We<br>stern Hospital<br>(TGH) | Centre for<br>Addition &<br>Mental Health<br>(CAM) | McMaster<br>University<br>(MCU) | University of<br>Calgary<br>(UCA) | University of<br>British<br>Columbia<br>(UBC) | Queens<br>University<br>(QNS) |
| --- | --- | --- | --- | --- | --- | --- |
| Scanner Model | GE 3T Signa HDxt | GE 3T Discovery MR750 | GE 3T Discovery MR750 | GE 3T Discovery MR750 | Philips 3T Intera | Siemens 3T TrioTim |
| Coil | GE 8HRBRAIN | GE 8HRBRAIN | GE HNS Head | GE HNS Head | SENSE-Head-8 | 12-channel head matrix coil |
| Software version | HD16.0_V02_1 131.a | DV24.0_R01_1 344.a | DV25.0_R02_1 549.b | DV25.0_R02_1 549.b | 3.2.3,3.2.3.1 | syngo MR B19 |
| TR (ms) | 7.5 <sup>[1]</sup> | 6.4 <sup>[2]</sup> | 6.4 <sup>[2]</sup> | 6.4 <sup>[2]</sup> | 6.57 | 1760 <sup>[3]</sup> |
| TE (ms) | 2.86 <sup>[4]</sup> | 2.8 <sup>[5]</sup> | 2.8 <sup>[5]</sup> | 2.8 <sup>[5]</sup> | 2.9 <sup>[6]</sup> | 2.2 <sup>[7]</sup> |
| TI (ms) | 450 | 450 | 450 | 450 | 950 | 950 <sup>[8]</sup> |
| Flip Angle (degree) | 15 | 15 | 15 | 15 | 8 | 15 |
| Pixel Bandwidth | 260 <sup>[9]</sup> | 260 <sup>[9]</sup> | 260 <sup>[10]</sup> | 260 <sup>[9]</sup> | 241 <sup>[11]</sup> | 199 |
| Matrix Dimension (pixels) | 240 x 240 <sup>[12]</sup> | 240 x 240 <sup>[12]</sup> | 240 x 240 <sup>[12]</sup> | 240 x 240 <sup>[12]</sup> | 240 x 240 <sup>[13]</sup> | 256 x 256 |
| Voxel Dimension (mm) | 1 x 1 x 1 | 1 x 1 x 1 | 1 x 1 x 1 | 1 x 1 x 1 <sup>[14]</sup> | 1 x 1 x 1 | 1 x 1 x 1 |
| Number of slices | 176 | 180 <sup>c1</sup> | 180 <sup>c1</sup> | 180 <sup>c1</sup> | 180 <sup>c2</sup> | 192 |
| Number of Subjects | 71 | 16 | 50 | 66 | 72 | 31 |

ONDRI:
| Site | Robarts Research Institute (ROB) | Sunnybrook Health Sciences (SBH) | St. Michael's Hospital (SMH) | The Ottawa Hospital (TOH) | Toronto Western Hospital (TWH) | Center for Addiction and Mental Health (CAM) |
| --- | --- | --- | --- | --- | --- | --- |
| Scanner | Siemens 3T Prisma fit | GE 3T Discovery MR750 | Siemens 3T Skyra | Siemens 3T Trio Tim | GE 3T Signa HDxt | GE 3T Discovery MR750 |
| TR (ms) | [2300:2300] | [8.156:8.156] | [2300:2300] | [2300:2300] | [6.9:7.3] | [6.652:6.652] |
| TE (ms) | [2.98:2.98] | [3.18:2.18] | [2.03:2.03] | [2.96:2.96] | [2.8:3.1] | [2.298:2.298] |
| TI (ms) | [900:900] | [400:400] | [900:900] | [900:900] | [400:400] | [400:400] |
| Flip Angle (degree) | [9:9] | [11:11] | [9:9] | [9:9] | [11:11] | [11:11] |
| Pixel Bandwidth | [240:240] | [244.141:244.141] | [240:240] | [240:240] | [244.141:244.141] | [244.141:244.141] |
| Matrix Dimension (pixels) | [256x256:256x256] | [256x256:256x256] | [256x256:256x256] | [256x256:256x256] | [256x256:256x256] | [256x256:256x256] |
| Voxel Dimension (mm) | [1x1x1 : 1x1x1] | [1x1x1 : 1x1x1] | [1x1x1 : 1x1x1] | [1x1x1 : 1x1x1] | [1x1x1 : 1x1x1] | [1x1x1 : 1x1x1] |
| Number of slices | [176:176] | [176:176] | [192:192] | [176:176] | [176:176] | [176:176] |
| Number of subjects | 17 | 29 | 12 | 38 | 30 | 13 |

## Appendix B Bootstrapped results of interrater reliability

The bootstrapped estimates (80% of the sample, repeated 100 times) of the IRR for the 3 raters for different combinations of the dataset and FreeSurfer versions are shown below:

|  | CANBIND v6.0 | ONDRI v6.0 | CANBIND v5.3 | ONDRI v5.3 |
| --- | --- | --- | --- | --- |
| ENQC | 0.279 (0.02) | 0.215 (0.033) | 0.361 (0.022) | 0.249 (0.026) |
| VisualQC | 0.635 (0.028) | 0.539 (0.046) | 0.586 (0.03) | 0.555 (0.041) |

## Appendix C: Distributions of derived features in erroneous and accurate ROIs

To help us better understand the differences between erroneous and accurate ROIs, we have visualized the distributions of derived features such as the cortical thickness between the subjects that were rated as erroneous and those that were not, for each Freesurfer label separately. They are shown in the plot below for the 12 most erroneous ROIs from the CANBIND dataset processed with FS v6.0. Distributions colored with green are from ROIs rated as accurate, and those colored with blue and red are from the erroneous ROIs from the healthy and disease cohorts respectively. It must be noted that we didn’t collect the exact coordinates of the errors in each FS label and are visualizing the distributions of thickness of the entire label from many thousands of vertices in each panel. Such massive distribution has the potential to drown any subtle differences from the exact location of erroneous vertices.

A clear pattern of errors we can see in these visualizations (shown below) are the peaks at 0mm (considered erroneous) for the labels entorhinal (Row 1 Col 3), parahippocampal (R2:C4) and temporal pole (R3:C4). While we do see a green peak (although smaller, from some fraction of subjects for the ROIs rated as not-erroneous), this is likely coming from slices not reviewed or missed by the quality rater, and serves as another reminder of how the complex the review process is and how tedious proper QC can be. However, we do notice much larger peaks at 0mm for the erroneous groups, which implies we were able to catch those errors with our QC protocol. We also see a noticeable difference in the shape of the no-error vs. error distributions in the panel corresponding temporal pole (R3:C4).

As noted before, given we are visualizing the distributions of thickness of the entire label from many thousands of vertices, we may be drowning any subtle differences, from the typical parcellation errors we notice in FS. When big global segmentation failures do happen, it can result in the 0mm peaks as identified earlier.

As the overlap of distributions is pretty clear, we don’t think it’s necessary to any statistics to show they do not significantly differ from each other. However, we included them (along with corresponding versions for sulcal depth and curvature), in the revised version for improved readability for the community.

**Fig C1:**
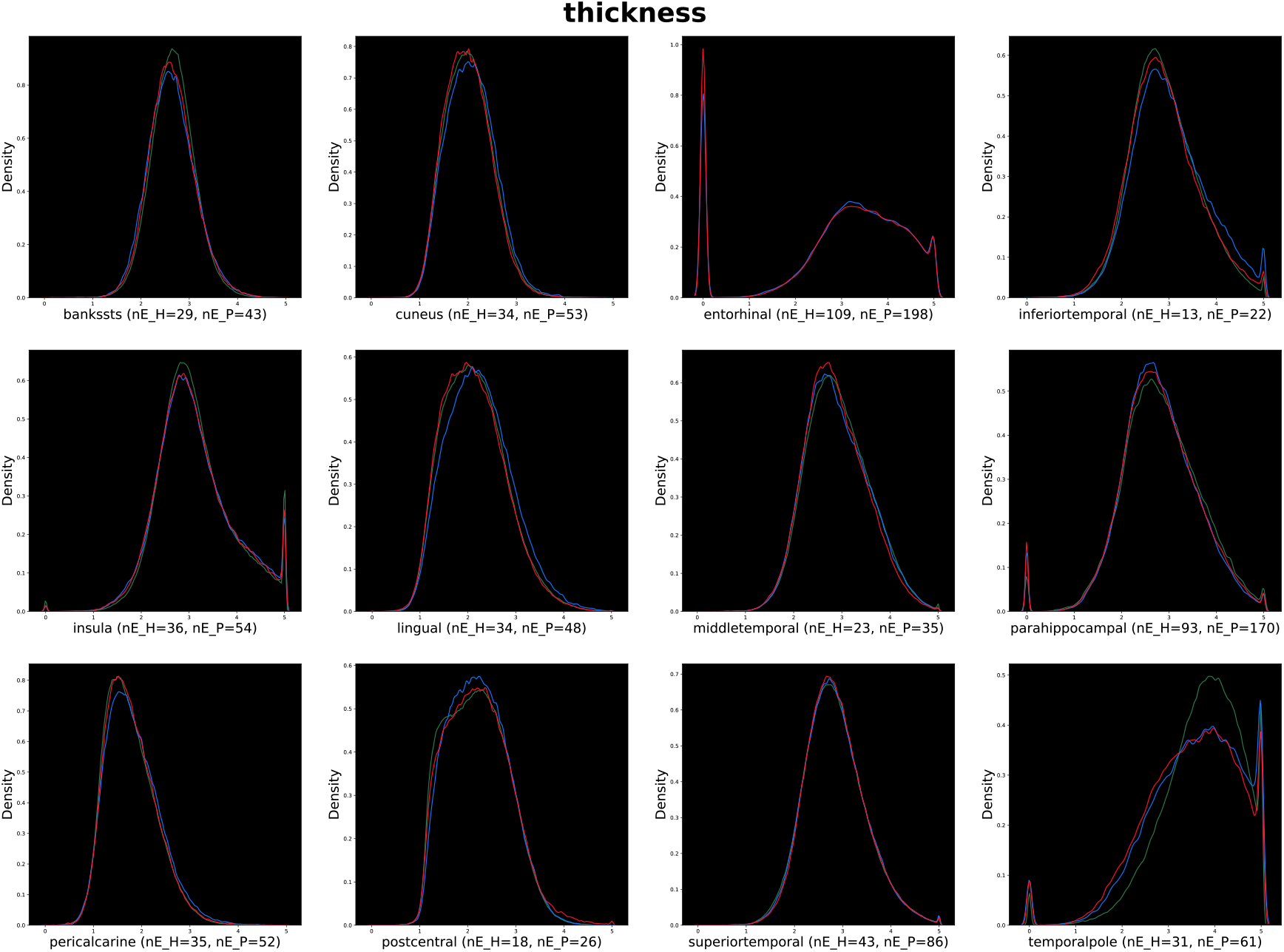
Distributions of vertex-wise cortical thickness in different ROIs grouped as erroneous or not (and further subdivided by healthy vs. patient). The x-axis refers to the thickness values, that are non-negative, with a typical average value of 2.5mm and a typical maximum value of about 5mm. The y-axis refers to the fraction of the ROIs at a particular value. The name of the ROI is noted in each panel’s x-axis label along with the number of erroneous subjects in each category.

**Fig C2:**
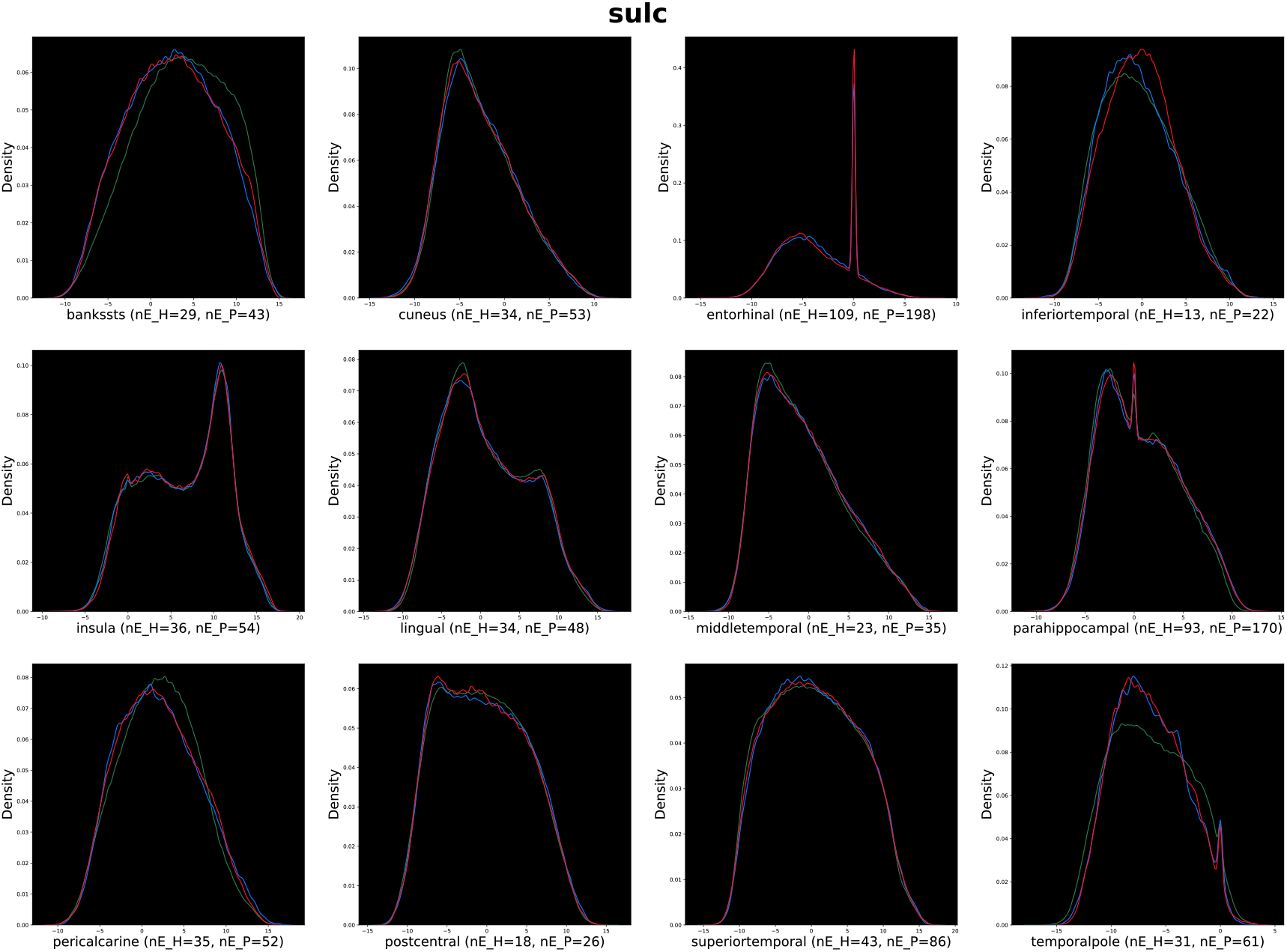
Distributions of vertex-wise sulcal depth in different ROIs rated and grouped as erroneous or not (and further subdivided by healthy vs. patient). The x-axis refers to the sulcal depth values, whose range includes negative values, whereas the y-axis refers to the fraction of the ROIs at a particular value. The name of the ROI is noted in each panel’s x-axis label along with the number of erroneous subjects in each category.

**Fig C3:**
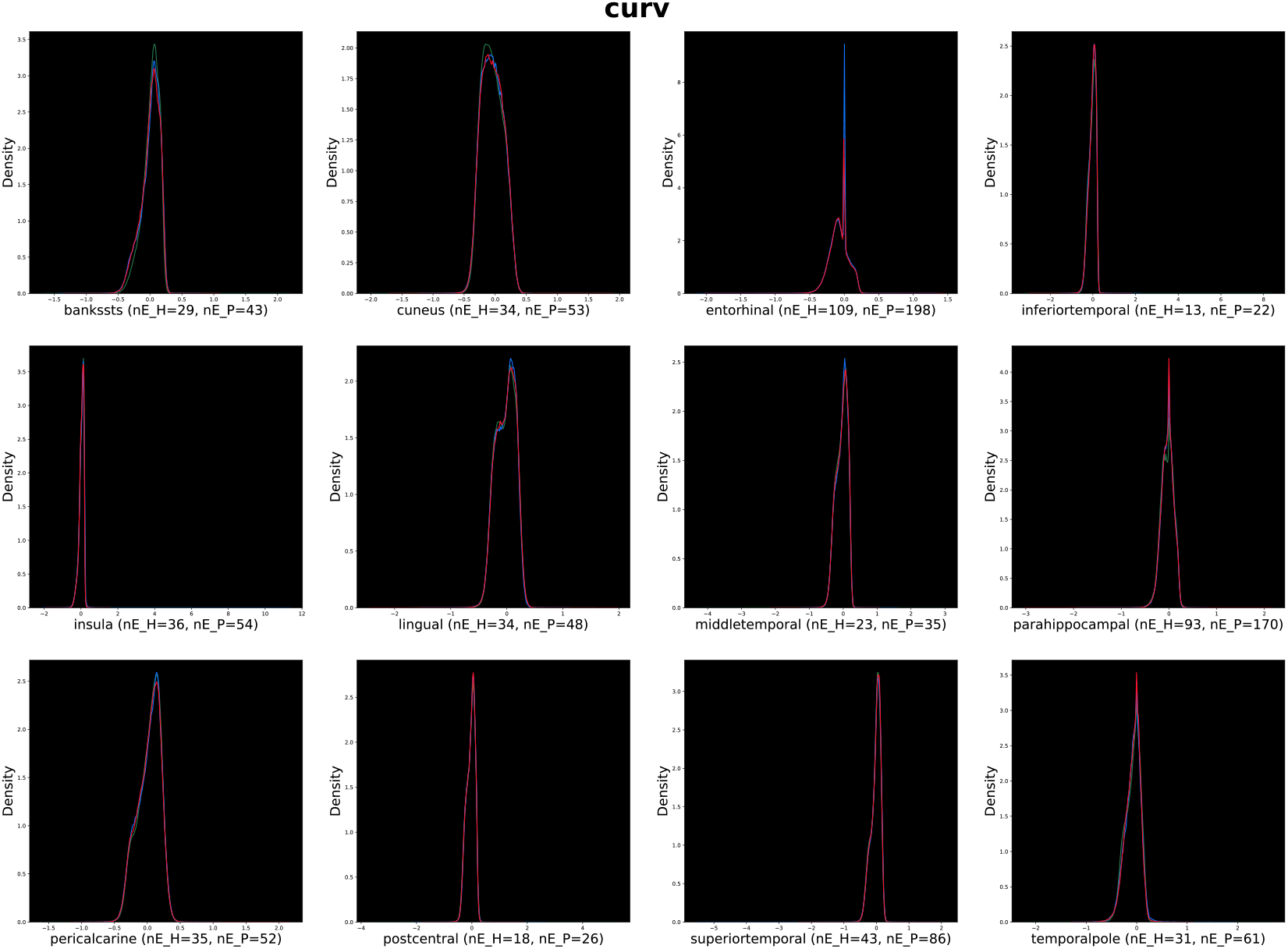
Distributions of vertex-wise curvature in different ROIs grouped as erroneous or not (and further subdivided by healthy vs. patient). The x-axis refers to the curvature values, whose range can include negative values, whereas the y-axis refers to the fraction of the ROIs at a particular value. The name of the ROI is noted in each panel’s x-axis label along with the number of erroneous subjects in each category.

## Acknowledgements

We like to thank the Indoc research team for their data management support. support. We would like to acknowledge the individuals and organizations that have made Data used for this research available including the Canadian Biomarker Integration Network in Depression (CAN-BIND), the Ontario Neurodegenerative Disease Research Initiative (ONDRI), the Ontario Brain Institute (OBI), the Canadian Open Neuroscience Platform (CONP), the Brain-CODE platform, and the Government of Ontario. The authors would like to acknowledge the ONDRI Founding Authors: Robert Bartha, Sandra E. Black, Michael Borrie, Dale Corbett, Elizabeth Finger, Morris Freedman, Barry Greenberg, David A. Grimes, Robert A. Hegele, Chris Hudson, Anthony E. Lang, Mario Masellis, William E. McIlroy, Paula

M. McLaughlin, Manuel Montero-Odasso, David G. Munoz, Douglas P. Munoz, J. B. Orange, Michael J. Strong, Stephen C. Strother, Richard H. Swartz, Sean Symons, Maria Carmela Tartaglia, Angela Troyer, and Lorne Zinman. The authors would also like to acknowledge the CAN-BIND Investigator Team: www.canbind.ca/our-team.

CAN-BIND is an Integrated Discovery Program carried out in partnership with, and financial support from, the Ontario Brain Institute, an independent non-profit corporation, funded partially by the Ontario government. The opinions, results and conclusions are those of the authors and no endorsement by the Ontario Brain Institute is intended or should be inferred. All study medications were independently purchased at wholesale market values.

The opinions, results, and conclusions are those of the authors and no endorsement by the Ontario Brain Institute is intended or should be inferred.

## Ethics Statement

All recruitment sites adopted a standardized Participant Agreement with the OBI to enable the transfer of data in accordance with the Governance Policy of OBI as well as the local institutional and/or ethical policies. Written and informed parental consent was obtained for all participants under the age of 16. The patients/participants provided their written informed consent to participate in this study. Written informed consent was obtained from the individual(s) for the publication of any potentially identifiable images or data included in this article.

## Footnotes

1 Please refer to the VisualQC manual for illustrations of the two protocols at URL: https://github.com/raamana/visualqc/blob/master/docs/VisualQC_TrainingManual_v1p4.pdf

2 CNR is computed as (`Mean(WM)-Mean(GM)`) / `sqrt` `((Var(WM)+Var(GM)))`, where all data used to compute means and variances are intensity values in WM/GM.

3 URL: https://github.com/raamana/visualqc/blob/master/docs/VisualQC_TrainingManual_v1p4.pdf

## References

Alfaro-Almagro, Fidel, Mark Jenkinson, Neal K Bangerter, Jesper L R Andersson, Ludovica Griffanti, Gwenaelle Douaud, Stamatios N Sotiropoulos, et al. 2018. “Image Processing and Quality Control for the First 10,000 Brain Imaging Datasets from UK Biobank.” NeuroImage 166 (February): 400–424. https://doi.org/10.1016/j.neuroimage.2017.10.034.

Backhausen, Lea L, Megan M Herting, Judith Buse, Veit Roessner, Michael N Smolka, and Nora C Vetter. 2016. “Quality Control of Structural MRI Images Applied Using FreeSurfer—A Hands-On Workflow to Rate Motion Artifacts.” Frontiers in Neuroscience 10 (January): 2385. https://doi.org/10.3389/fnins.2016.00558.

ENIGMA Consortium, The. 2017. “ENIGMA Imaging Protocols.” 2017. http://enigma.ini.usc.edu/protocols/imaging-protocols/.

Esteban, Oscar, Daniel Birman, Marie Schaer, Oluwasanmi O Koyejo, Russell A Poldrack, and Krzysztof J Gorgolewski. 2017. “MRIQC: Advancing the Automatic Prediction of Image Quality in MRI from Unseen Sites.” edited by Boris C Bernhardt. PLoS ONE 12 (9): e0184661. https://doi.org/10.1371/journal.pone.0184661.

Farhan, Sali M. K., Robert Bartha, Sandra E. Black, Dale Corbett, Elizabeth Finger, Morris Freedman, Barry Greenberg, et al. 2017. “The Ontario Neurodegenerative Disease Research Initiative (ONDRI).” Canadian Journal of Neurological Sciences / Journal Canadien Des Sciences Neurologiques 44 (2): 196–202. https://doi.org/10.1017/cjn.2016.415.

Fischl, Bruce. 2012. “FreeSurfer.” NeuroImage 62 (2): 774–81. https://doi.org/10.1016/j.neuroimage.2012.01.021.

Fleiss, Joseph L. 1971. “Measuring Nominal Scale Agreement among Many Raters.” Psychological Bulletin 76 (5): 378–82. https://doi.org/10.1037/h0031619.

Freesurfer Team. 2017. “Official Troubleshooting Guide.” 2017. https://surfer.nmr.mgh.harvard.edu/fswiki/FsTutorial/TroubleshootingData.

Gedamu, Elias L., D.L. Collins, and Douglas L. Arnold. 2008. “Automated Quality Control of Brain MR Images.” Journal of Magnetic Resonance Imaging 28 (2): 308–19. https://doi.org/10.1002/jmri.21434.

Keshavan, Anisha, Esha Datta, Ian M. McDonough, Christopher R. Madan, Kesshi Jordan, and Roland G. Henry. 2018. “Mindcontrol: A Web Application for Brain Segmentation Quality Control.” NeuroImage 170 (April): 365–72. https://doi.org/10.1016/j.neuroimage.2017.03.055.

Klapwijk, Eduard T., Ferdi van de Kamp, Mara van der Meulen, Sabine Peters, and Lara M. Wierenga. 2019. “Qoala-T: A Supervised-Learning Tool for Quality Control of FreeSurfer Segmented MRI Data.” NeuroImage 189 (April): 116–29. https://doi.org/10.1016/j.neuroimage.2019.01.014.

Lam, Raymond W., Roumen Milev, Susan Rotzinger, Ana C. Andreazza, Pierre Blier, Colleen Brenner, Zafiris J. Daskalakis, et al. 2016. “Discovering Biomarkers for Antidepressant Response: Protocol from the Canadian Biomarker Integration Network in Depression (CAN-BIND) and Clinical Characteristics of the First Patient Cohort.” BMC Psychiatry 16 (1). https://doi.org/10.1186/s12888-016-0785-x.

MacQueen, Glenda M., Stefanie Hassel, CAN-BIND Investigator Team, Stephen R. Arnott, Jean Addington, Christopher R. Bowie, Signe L. Bray, et al. 2019. “The Canadian Biomarker Integration Network in Depression (CAN-BIND): Magnetic Resonance Imaging Protocols.” Journal of Psychiatry and Neuroscience 44 (4): 223–36. https://doi.org/10.1503/jpn.180036.

Marcus, Daniel S, Michael P Harms, Abraham Z Snyder, Mark Jenkinson, J Anthony Wilson, Matthew F Glasser, Deanna M Barch, et al. 2013. “Human Connectome Project Informatics: Quality Control, Database Services, and Data Visualization.” NeuroImage 80 (October): 202–19. https://doi.org/10.1016/j.neuroimage.2013.05.077.

Mortamet, Bénédicte, Matt A Bernstein, Clifford R Jack, Jeffrey L Gunter, Chadwick Ward, Paula J Britson, Reto Meuli, Jean-Philippe Thiran, and Gunnar Krueger. 2009. “Automatic Quality Assessment in Structural Brain Magnetic Resonance Imaging.” Magnetic Resonance in Medicine 62 (2): 365–72. https://doi.org/10.1002/mrm.21992.

Pizarro, Ricardo A, Xi Cheng, Alan Barnett, Herve Lemaitre, Beth A Verchinski, Aaron L Goldman, Ena Xiao, et al. 2016. “Automated Quality Assessment of Structural Magnetic Resonance Brain Images Based on a Supervised Machine Learning Algorithm.” Frontiers in Neuroinformatics 10 (December): 805. https://doi.org/10.3389/fninf.2016.00052.

Raamana, Pradeep Reddy. 2018. “VisualQC: Assistive Tools For Easy And Rigorous Quality Control Of Neuroimaging Data,” April. https://doi.org/10.5281/ZENODO.1211365.

Randolph, Justus J. 2005. “Free-Marginal Multirater Kappa (Multirater K [Free]): An Alternative to Fleiss’ Fixed-Marginal Multirater Kappa.” In Joensuu Learning and Instruction Symposium. https://eric.ed.gov/?id=ED490661.

Rosen, Adon F G, David R Roalf, Kosha Ruparel, Jason Blake, Kevin Seelaus, Lakshmi P Villa, Rastko Ciric, et al. 2017. “Quantitative Assessment of Structural Image Quality.” NeuroImage 169 (December): 407–18. https://doi.org/10.1016/j.neuroimage.2017.12.059.

Scott, Christopher J.M., Stephen R. Arnott, Aditi Chemparathy, Fan Dong, Igor Solovey, Tom Gee, Tanya Schmah, et al. 2020. “An Overview of the Quality Assurance and Quality Control of Magnetic Resonance Imaging Data for the Ontario Neurodegenerative Disease Research Initiative (ONDRI): Pipeline Development and Neuroinformatics.” Preprint. Neuroscience. https://doi.org/10.1101/2020.01.10.896415.

Seabold, Skipper, and Josef Perktold. 2010. Statsmodels: Econometric and Statistical Modeling with Python. 9th Python in Science Conference.

Shehzad, Zarrar, Giavasis Steven, Li Qingyang, Benhajali Yassine, Yan Chaogan, Yang Zhen, Milham Michael, Bellec Pierre, and Craddock Cameron. 2015. “The Preprocessed Connectomes Project Quality Assessment Protocol - a Resource for Measuring the Quality of MRI Data.” Frontiers in Neuroscience 9. https://doi.org/10.3389/conf.fnins.2015.91.00047.

SIG, niQC. 2019. “Neuroimaging Quality Control (NiQC) Special Interest Group at the INCF,” 2019. https://incf.github.io/niQC/tools.

Thompson, Paul M., Neda Jahanshad, Christopher R. K. Ching, Lauren E. Salminen, Sophia I. Thomopoulos, Joanna Bright, Bernhard T. Baune, et al. 2020. “ENIGMA and Global Neuroscience: A Decade of Large-Scale Studies of the Brain in Health and Disease across More than 40 Countries.” Translational Psychiatry 10 (1). https://doi.org/10.1038/s41398-020-0705-1.

White, Tonya, Philip R. Jansen, Ryan L. Muetzel, Gustavo Sudre, Hanan El Marroun, Henning Tiemeier, Anqi Qiu, Philip Shaw, Andrew M. Michael, and Frank C. Verhulst. 2018. “Automated Quality Assessment of Structural Magnetic Resonance Images in Children: Comparison with Visual Inspection and Surface-Based Reconstruction.” Human Brain Mapping 39 (3): 1218–31. https://doi.org/10.1002/hbm.23911.

Woodard, Jeffrey P., and Monica P. Carley-Spencer. 2006. “No-Reference Image Quality Metrics for Structural MRI.” Neuroinformatics 4 (3): 243–62. https://doi.org/10.1385/NI:4:3:243.

